## Supplemental Tables and Figures for "KMD clustering: Robust general-purpose clustering of biological data"

**Supplementary information**

**Supplementary Tables**

| | **nested circles** (1000 objects, 2 clusters) | | | **half moons** (1000 objects, 2 clusters) | | | **globular clusters** (1000 objects, 3 clusters) | | | **anisotropic clusters** (1000 objects, 3 clusters**)** | | |
|---|---|---|---|---|---|---|---|---|---|---|---|---|
| | Accuracy | NMI | ARI | Accuracy | NMI | ARI | Accuracy | NMI | ARI | Accuracy | NMI | ARI |
| Spectral | 1 | 1 | 1 | 1 | 1 | 1 | 0.962 | 0.849 | 0.891 | 0.726 | 0.838 | 0.853 |
| Average Linkage | 0.661 | 0.230 | 0.103 | 0.897 | 0.612 | 0.63 | 0.697 | 0.63 | 0.527 | 0.428 | 0.694 | 0.699 |
| Single Linkage | 1 | 1 | 1 | 1 | 1 | 1 | 0.335 | 0.012 | 0 | 0.350 | 0.06 | 0 |
| DBSCAN | 0.820 | 0.720 | 0.733 | 1 | 1 | 1 | 0.970 | 0.911 | 0.941 | 0.984 | 0.968 | 0.977 |
| HDBSCAN | 1 | 1 | 1 | 1 | 1 | 1 | 0.984 | 0.934 | 0.96 | 1 | 1 | 1 |
| Gaussian Mixture | 0.512 | 0 | 0 | 0.851 | 0.393 | 0.492 | 0.966 | 0.86 | 0.901 | 0.784 | 0.994 | 0.997 |
| KMD | 1 | 1 | 1 | 1 | 1 | 1 | 0.977 | 0.847 | 0.888 | 0.999 | 0.974 | 0.985 |
| KMD core | 1 | 1 | 1 | 1 | 1 | 1 | 0.983 | 0.876 | 0.919 | 0.994 | 0.994 | 0.997 |

**Table S1. Evaluation on simulated datasets.** Performance metrics on scikit-learn datasets including accuracy, Normalized Mutual Information Adjusted Rand Index.

| | **nested circles** (1000 objects, 2 clusters) | | | **half moons** (1000 objects, 2 clusters) | | | **globular clusters** (1000 objects, 3 clusters) | | | **anisotropic clusters** (1000 objects, 3 clusters**)** | | |
|---|---|---|---|---|---|---|---|---|---|---|---|---|
| | Accuracy | NMI | ARI | Accuracy | NMI | ARI | Accuracy | NMI | ARI | Accuracy | NMI | ARI |
| Spectral | 0.711 | 0.289 | 0.177 | 0.853 | 0.403 | 0.5 | 0.899 | 0.665 | 0.726 | 0.91 | 0.741 | 0.76 |
| Average Linkage | 0.687 | 0.237 | 0.139 | 0.839 | 0.41 | 0.459 | 0.632 | 0.514 | 0.428 | 0.668 | 0.722 | 0.568 |
| Single Linkage | 0.501 | 0.002 | 0 | 0.501 | 0.002 | 0 | 0.335 | 0.004 | 0 | 0.335 | 0.004 | 0 |
| DBSCAN | 0.667 | 0.536 | 0.721 | 0.494 | 0.352 | 0 | 0.342 | 0.033 | 0 | 0.67 | 0.724 | 0.567 |
| HDBSCAN | 0 | 0 | 0 | 0.941 | 0.678 | 0.778 | 0.924 | 0.754 | 0.794 | 0.667 | 0.736 | 0.569 |
| Gaussian Mixture | 0.552 | 0.009 | 0.011 | 0.834 | 0.353 | 0.457 | 0.923 | 0.727 | 0.784 | 0.998 | 0.976 | 0.988 |
| KMD | 0.990 | 0.922 | 0.960 | 0.915 | 0.581 | 0.689 | 0.914 | 0.717 | 0.763 | 0.971 | 0.881 | 0.916 |
| KMD core | 0.992 | 0.932 | 0.967 | 0.929 | 0.631 | 0.736 | 0.916 | 0.726 | 0.772 | 0.995 | 0.973 | 0.984 |

**Table S2. Evaluation on simulated noisy datasets.** Performance metrics on scikit-learn datasets (with high noise added) including accuracy, Normalized Mutual Information Adjusted Rand Index.

| | **nested circles** (1000 objects, 2 clusters) | | | **half moons** (1000 objects, 2 clusters) | | | **globular clusters** (1000 objects, 3 clusters) | | | **anisotropic clusters** (1000 objects, 3 clusters**)** | | |
|---|---|---|---|---|---|---|---|---|---|---|---|---|
| | Accuracy | NMI | ARI | Accuracy | NMI | ARI | Accuracy | NMI | ARI | Accuracy | NMI | ARI |
| Average Linkage | 0.687 | 0.237 | 0.139 | 0.839 | 0.41 | 0.459 | 0.632 | 0.514 | 0.428 | 0.668 | 0.722 | 0.568 |
| Single Linkage | 0.501 | 0.002 | 0 | 0.501 | 0.002 | 0 | 0.335 | 0.004 | 0 | 0.335 | 0.004 | 0 |
| Complete Linkage | 0.804 | 0.364 | 0.369 | 0.834 | 0.367 | 0.446 | 0.863 | 0.621 | 0.643 | 0.582 | 0.471 | 0.376 |
| Ward Linkage | 0.662 | 0.184 | 0.104 | 0.756 | 0.205 | 0.261 | 0.903 | 0.694 | 0.729 | 0.844 | 0.741 | 0.652 |
| Minimax Linkage | 0.699 | 0.253 | 0.158 | 0.742 | 0.205 | 0.234 | 0.890 | 0.660 | 0.698 | 0.680 | 0.598 | 0.475 |
| KMD | 0.990 | 0.922 | 0.960 | 0.915 | 0.581 | 0.689 | 0.914 | 0.717 | 0.763 | 0.971 | 0.881 | 0.916 |
| KMD core | 0.992 | 0.932 | 0.967 | 0.929 | 0.631 | 0.736 | 0.916 | 0.726 | 0.772 | 0.995 | 0.973 | 0.984 |

**Table S3. Evaluation of different linkage methods on simulated noisy datasets.** Performance metrics on scikit-learn datasets (with high noise added) including accuracy, Normalized Mutual Information Adjusted Rand Index.

| | **nested circles** (1000 objects, 2 clusters) | | | **half moons** (1000 objects, 2 clusters) | | | **globular clusters** (1000 objects, 3 clusters) | | | **anisotropic clusters** (1000 objects, 3 clusters**)** | | |
|---|---|---|---|---|---|---|---|---|---|---|---|---|
| | Acc. | NMI | ARI | Acc. | NMI | ARI | Acc. | NMI | ARI | Acc. | NMI | ARI |
| Louvain | 0.19 | 0.387 | 0.142 | 0.187 | 0.305 | 0.131 | 0.0347 | 0.594 | 0.313 | 0.296 | 0.48 | 0.247 |
| Leiden | 0.138 | 0.350 | 0.100 | 0.155 | 0.268 | 0.089 | 0.232 | 0.535 | 0.218 | 0.213 | 0.442 | 0.176 |
| Average SCCAF | 0.56 | 0.348 | 0.292 | 0.553 | 0.152 | 0.068 | 0.675 | 0.642 | 0.522 | 0.460 | 0.206 | 0.150 |
| KMD | 0.990 | 0.922 | 0.960 | 0.915 | 0.581 | 0.689 | 0.914 | 0.717 | 0.763 | 0.971 | 0.881 | 0.916 |
| KMD core | 0.992 | 0.932 | 0.967 | 0.929 | 0.631 | 0.736 | 0.916 | 0.726 | 0.772 | 0.995 | 0.973 | 0.984 |

**Table S4. Evaluation of different single-cell clustering methods on simulated noisy datasets.** Evaluation of methods by accuracy, Normalized Mutual Information, Adjusted Rand Index. Comparison of clustering algorithm performance on standard scikit-learn simulated datasets with high noise added. SCCAF was run 10 times to offset the effect of randomness.

| | Levine15_13 (20000 out of 167044 cells, 13 markers, 24 clusters) | | | Levine15_32 (20000 out of 265627 cells, 32 markers, 14 clusters) | | | Samusik16 (20000 out of 86864 cells, 44 markers, 24 clusters) | | |
|---|---|---|---|---|---|---|---|---|---|
| | accuracy | NMI | ARI | accuracy | NMI | ARI | accuracy | NMI | ARI |
| kmeans | 0.5587+-0.0303 | 0.7071+-0.0139 | 0.5642+-0.0367 | 0.5702+-0.0550 | 0.7252+-0.0213 | 0.6169+-0.0755 | 0.4819+-0.0585 | 0.6506+-0.0201 | 0.4628+-0.0512 |
| Xshift | 0.7824+-0.0216 | 0.7849+-0.0205 | 0.7722+-0.0298 | 0.8706+-0.0510 | 0.8646+-0.0405 | 0.8958+-0.04278 | 0.9100+-0.0230 | 0.8702+-0.0063 | 0.8890+-0.0281 |
| DEPECHE | 0.6918+-0.0141 | 0.6679+-0.0094 | 0.6559+-0.0107 | 0.8922+-0.0023 | 0.8416+-0.0030 | 0.9273+-0.0031 | 0.8253+-0.0067 | 0.7243+-0.0059 | 0.8251+-0.0079 |
| Accense | 0.5591 +-0.1203 | 0.7224 +-0.061 | 0.5911 +-0.129 | 0.5218+-0.0947 | 0.6644+-0.0590 | 0.4811+-0.1048 | 0.5935+-0.0714 | 0.7279+-0.0383 | 0.5664+-0.0767 |
| FlowSOM | 0.8436+-0.0272 | 0.8443+-0.0143 | 0.8579+-0.0209 | 0.8646+-0.0903 | 0.8725+-0.0596 | 0.8734+-0.1195 | 0.6331+-0.0666 | 0.6599+-0.0460 | 0.6188+-0.0833 |
| PhenoGraph | 0.9180+-0.0014 | 0.8827+-0.0021 | 0.9268+-0.0032 | 0.6598+-0.0412 | 0.7607+-0.0220 | 0.6719+-0.0368 | 0.9235+-0.0423 | 0.8996+-0.0257 | 0.9248+-0.0524 |
| KMD | 0.8117+-0.0448 | 0.8073+-0.0149 | 0.7951+-0.0363 | 0.9479+-0.0022 | 0.9435+-0.0035 | 0.9706+-0.0016 | 0.9221+-0.0058 | 0.8873+-0.0044 | 0.9089+-0.0034 |

**Table S5. Evaluation on mass cytometry data.** Average and standard deviation of performance metrics on mass cytometry datasets including accuracy, Normalized Mutual Information Adjusted Rand Index.

| | **Lawlor17**<br><br>(638 cells,19927 genes, 8 clusters) | | | **Zeisel15**<br><br>(2361 cells, 2000 genes, 7 clusters) | | | **Li17**<br><br>(561 cells, 25083 genes, 7 clusters) | | |
|---|---|---|---|---|---|---|---|---|---|
| | accuracy | NMI | ARI | accuracy | NMI | ARI | accuracy | NMI | ARI |
| Louvain | 0.674 | 0.699 | 0.576 | 0.623 | 0.728 | 0.496 | 0.729 | 0.764 | 0.592 |
| Leiden | 0.686 | 0.723 | 0.583 | 0.52 | 0.683 | 0.400 | 0.713 | 0.783 | 0.586 |
| Average SCCAF | 0.808 | 0.76 | 0.769 | 0.711 | 0.731 | 0.586 | 0.582 | 0.5 | 0.341 |
| Seurat | 0.657 | 0.678 | 0.658 | 0.611 | 0.716 | 0.488 | 0.715 | 0.687 | 0.544 |
| KMD | 0.893 | 0.790 | 0.831 | 0.738 | 0.686 | 0.523 | 0.838 | 0.821 | 0.703 |

**Table S6. Evaluation on scRNA-seq datasets.** Performance metrics scRNA-Seq datasets including accuracy, Normalized Mutual Information Adjusted Rand Index.

| | Simulated scRNA-seq 150 objects, 3 clusters | | |
|---|---|---|---|
| | Accuracy | NMI | ARI |
| Louvain | 0.993 | 0.970 | 0.989 |
| Leiden | 0.987 | 0.950 | 0.960 |
| KMD | 0.987 | 0.940 | 0.960 |

**Table S7. Evaluation of different single-cell clustering methods on simulated scRNA-seq dataset.** Performance metrics on simulated scRNA-Seq dataset including accuracy, Normalized Mutual Information Adjusted Rand Index.

### Supplementary figures

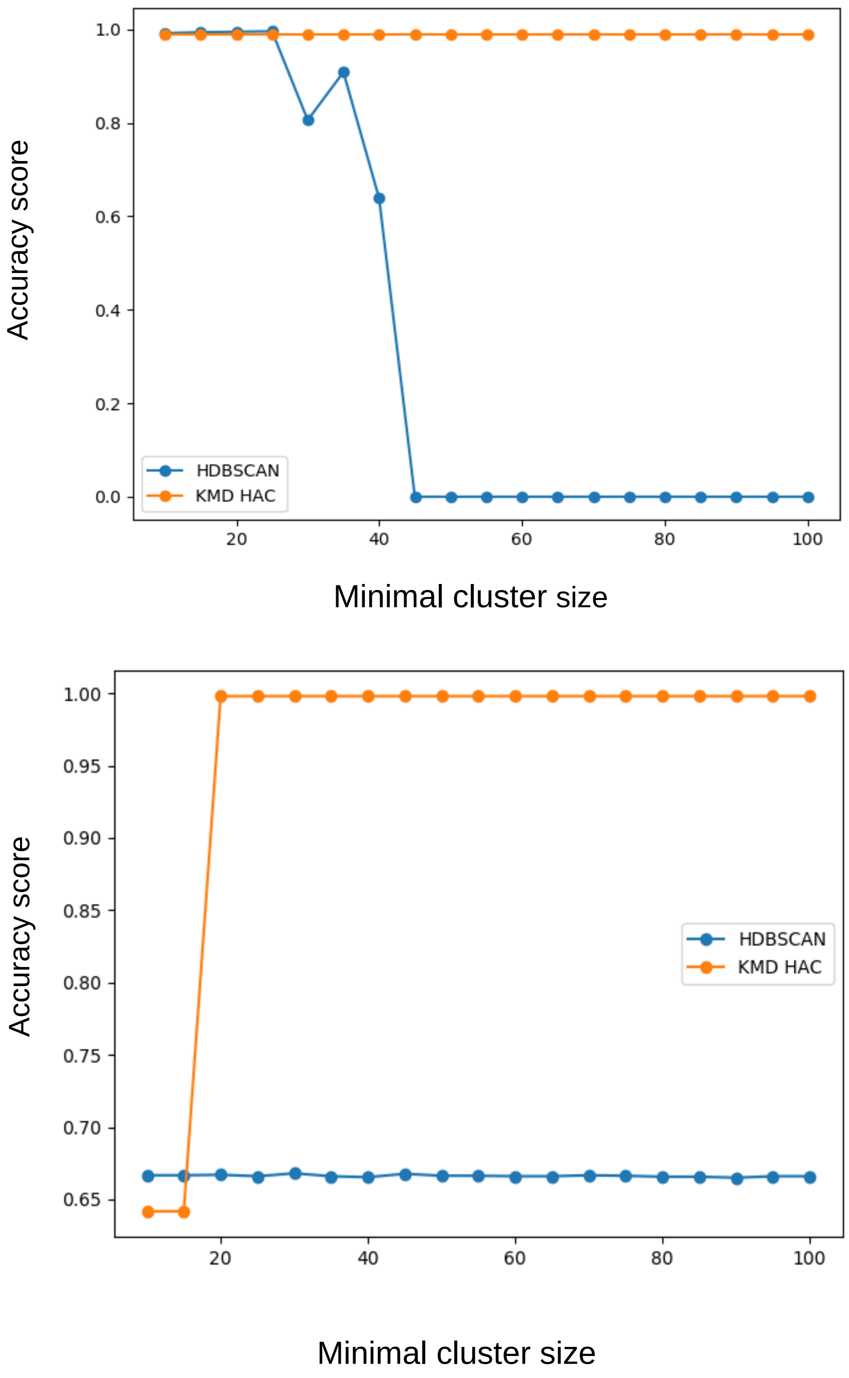


**Figure S1. Effect of minimal cluster size of clustering accuracy.** Clustering accuracy of KMD clustering and HDBSCAN on the high-noise nested circles (top) and high noise anisotropic clusters (bottom) datasets, shown for a range of minimal cluster size values.

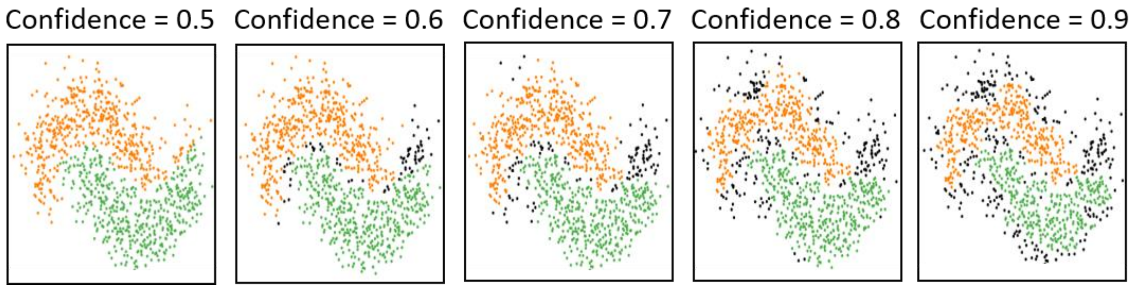


**Figure S2. Effect of outlier assignment confidence threshold.** Outlier cluster assignment at various confidence thresholds.

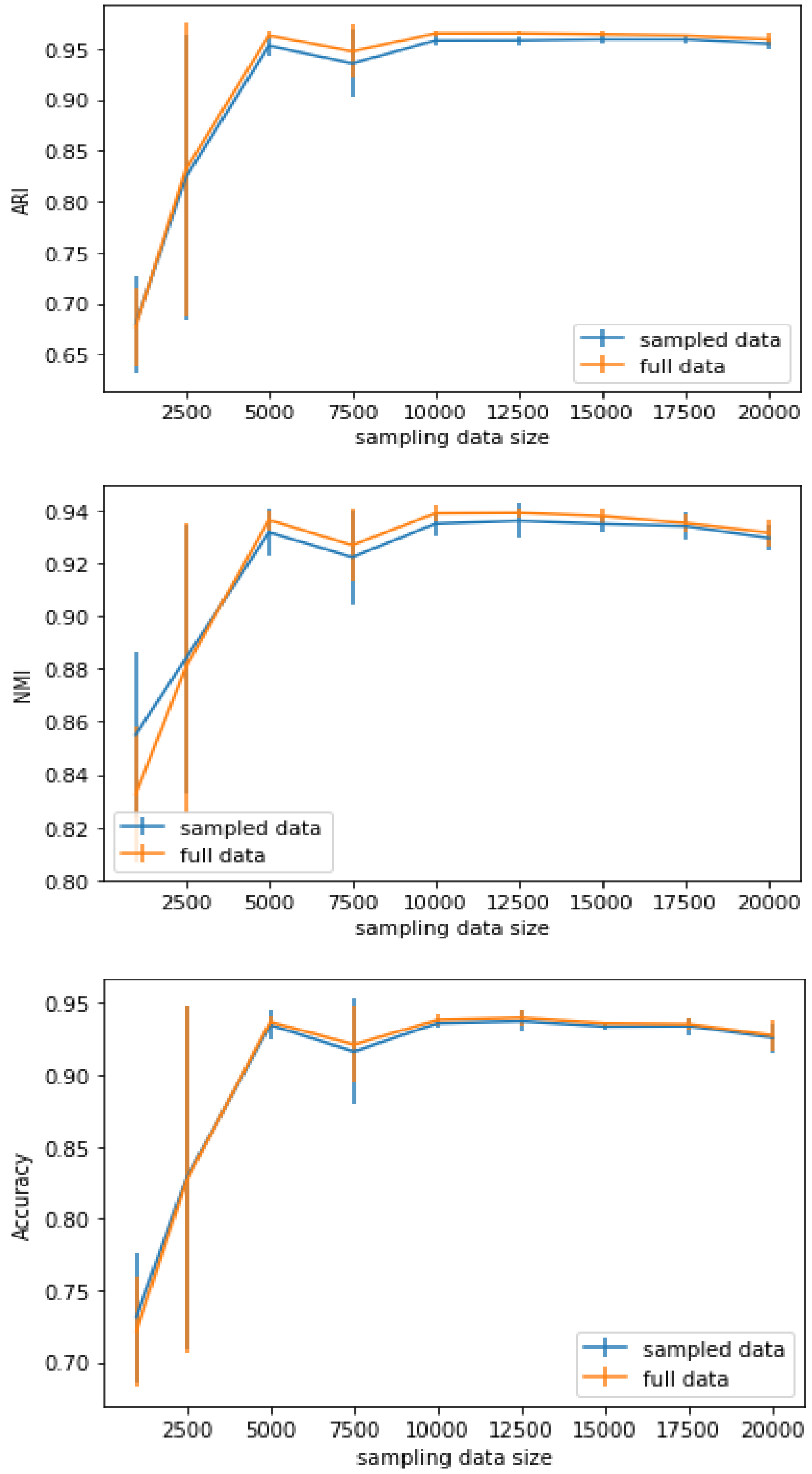


**Figure S3. Clustering performance using sampling-based approach.** Sampling-based KMD clustering was performed on the full Levine32dim dataset by randomly sampling a subset of cells of the specified sizes (1000-20000). ARI, NMI and accuracy were evaluated both for the sampled subset and for the entire data (i.e. the sampled subset which for the core clusters as well as the rest of the cells which were assigned based on the core clusters).

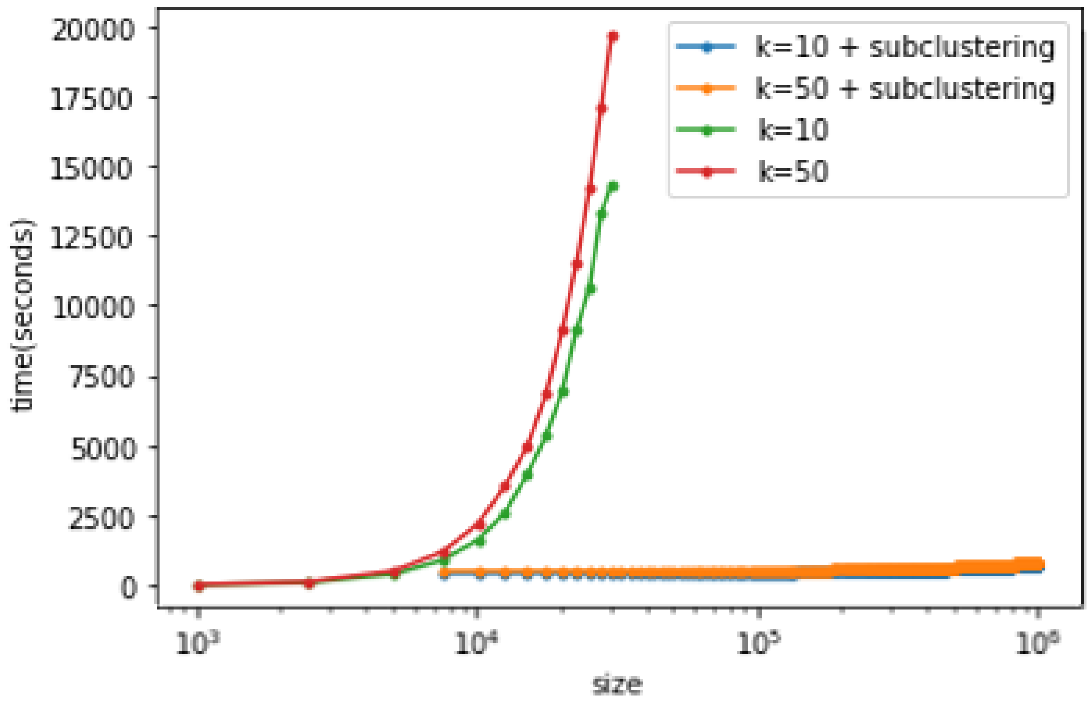


**Figure S4. KMD clustering run time on large dataset.** KMD clustering runtime was evaluated on random samples taken from the simZeisel15 dataset which contains 1M objects. Two different k values (10 and 50) are shown. Full KMD clustering was performed for datasets of up to 20000 objects. Sampling-based KMD clustering (marked “subclustering”) was performed on datasets of up to 1M objects, while the sample size was kept constant at 5000 objects.
